# Test-retest reliability of TMS-evoked potentials over fMRI-based definitions of non-motor cortical targets

**DOI:** 10.1101/2024.12.20.629675

**Authors:** Ceyda Sayalı, Juha Gogulski, Ida Granö, Pantelis Lioumis, Frederick S. Barrett

## Abstract

**Objective:** This study assessed the test-retest reliability of TMS-evoked potentials (TEPs) across two cortical regions—dorsolateral prefrontal cortex (DLPFC), and angular gyrus— in comparison to motor cortex (M1), using individualized and literature-based targeting approaches. The study compared the reliability of single-pulse TMS, short-interval intracortical inhibition (SICI), and long-interval intracortical inhibition (LICI) protocols to evaluate TEP consistency in these regions.

**Methods:** Seventeen healthy participants underwent two TMS-EEG sessions spaced by at least one week, with targets for DLPFC and angular gyrus identified using resting-state functional connectivity (RS) and Neurosynth-based functional overlays. Motor cortex was targeted using resting motor threshold (RMT). Early TEPs were quantified as peak-to-peak amplitude, in dBμV. Test-retest reliability of early TEPs was calculated using the concordance correlation coefficient (CCC) for each region and protocol.

**Results:** M1 demonstrated the highest TEP reliability (CCCmean = 0.59), while DLPFC (CCCmean = 0.40) and angular gyrus (CCCmean = 0.45) showed lower reliability, particularly for anterior DLPFC targets. Neurosynth-based DLPFC targets exhibited slightly higher CCC values (mean CCC = 0.57) compared to RS-based targets (mean CCC = 0.30), but the difference was not statistically significant. No significant differences in reliability were found across single pulse and paired pulse protocols. Lateral targets, DLPFC and angular gyrus, showed lower reliability in comparison to motor cortex which might have been caused by muscle artifacts.

**Conclusion:** While individualized functional targeting methods provide advantages in engaging specific brain networks, their reliability for TEP measurements remains lower than the RMT-based approach for motor cortex. Future studies should integrate neuroimaging-based targeting with real-time TEP monitoring to enhance reliability in non-motor regions. This approach could enhance the precision of TMS-EEG protocols, especially for clinical applications targeting cortical regions like the DLPFC and angular gyrus.

## Introduction

Transcranial magnetic stimulation (TMS) precisely manipulates brain activity, influencing behavior and cognition^1,2,3,4^. Various methods for determining stimulation targets have emerged, including anatomical landmarks, brain atlas coordinates, and individualized MRI-derived coordinates, the latter offering greater precision by accounting for unique brain anatomy^5,6,7,8,9,10,11,12^. Individualized approaches typically yield more reliable target engagement^13,14,15,16,17,18^.

fMRI-guided TMS targeting, particularly using resting-state connectivity, has shown promise for engaging regions like the dorsolateral prefrontal cortex (DLPFC), a key target in treating major depressive disorder (MDD^19^). Functional connectivity between the DLPFC and subgenual cingulate cortex predicts therapeutic outcomes^20^. Standard clinical targeting protocols (e.g., 5.5 cm anterior to the motor threshold site^21^) overlook substantial variability in DLPFC anatomy^22,23,24,25^. Individualized targeting has demonstrated stronger treatment outcomes with fewer participants than standardized methods^15^. Consequently, cognitive neuroscience increasingly relies on fMRI-based targeting to engage specific brain regions^26,27,28,29,30,31,32^.

TMS-EEG combines TMS with electrophysiological measurements, offering insights into brain activity, connectivity, and plasticity^33,34,35^. Its reproducibility is well-documented^36,37,38^ and valuable for studying disorders like MDD and schizophrenia^36,39,40,41,42,43,44,45,46,47,48,49,50,51,52,53,54,55,56,57^. However, TMS-EEG targeting predominantly relies on standardized methods like EEG electrode placement, which are prone to artifacts and variability, underscoring the need for precision^9,58,59,60,61,62,63,64,65,66,67^.

Although the test-retest reliability of non-motor regions such as the DLPFC has been investigated^36,66,68,69,70^, the effects of different targeting strategies remain unexplored. Furthermore, studies investigating test-retest reliability have utilized EEG cap placement to guide TMS targeting. This study compares the reliability of early TMS-evoked potentials (TEPs) across three brain regions—motor cortex, DLPFC, and angular gyrus—using resting-state fMRI and literature-derived targeting methods, using three TMS protocols (single pulse, SICI, LICI). The motor cortex, targeted via motor hotspot, serves as a reliability benchmark, while DLPFC and angular gyrus, key nodes in the frontoparietal (FPN) and default mode networks (DMN), respectively, are examined for their roles in cognition and mood regulation^71,72^. These comparisons provide insights into how targeting methods impact TEP reliability across distinct cortical regions.

## Materials and Methods

### Participants

Eighteen healthy participants participated in the study. One participant did not complete the second TMS-EEG session, and their data was excluded from the analysis. Seventeen participants (*Mean_age_* = 28, *SD_age_* = 6.28, 8 women and 9 men) completed all study procedures. Participants with psychiatric or neurological disorders (including moderate or severe depression), recent substance use disorder, cardiac events, pregnancy, contraindications for TMS or MRI, or a lifetime history of psychotropic medication use were excluded. The study’s experimental procedures were approved by the Johns Hopkins University Institutional Review Board. All participants provided written informed consent before the experiments.

### Design

After a baseline MRI acquisition, the study followed a repeated-measures design, with participants visiting the lab on two separate occasions separated by at least one week (**Figure 1A**). All TMS-EEG procedures were repeated at each visit. To ensure consistency, each session began at the same time of day for both visits. During each session, participants were instructed to keep their eyes open and remain still throughout the recordings. Active noise masking was applied using white noise mixed with click sounds^73^, delivered through sound-isolating (up to 29dB) in-ear earphones (Compy Foam Ear Canal Tips, Comply, Inc, North Oakdale, MN, USA). Before the experiment, the noise level was adjusted below a level of discomfort for the participant, but until the participant could no longer hear the TMS click at 100% maximum stimulator output (MSO) with the coil positioned 10 cm above the vertex. For parietal cortex stimulations, participants rested their chin on a chin rest to maintain a stable head position throughout the procedure. Participants were instructed to keep their gaze steady but were allowed to move their eyes freely.

**Figure 1.**
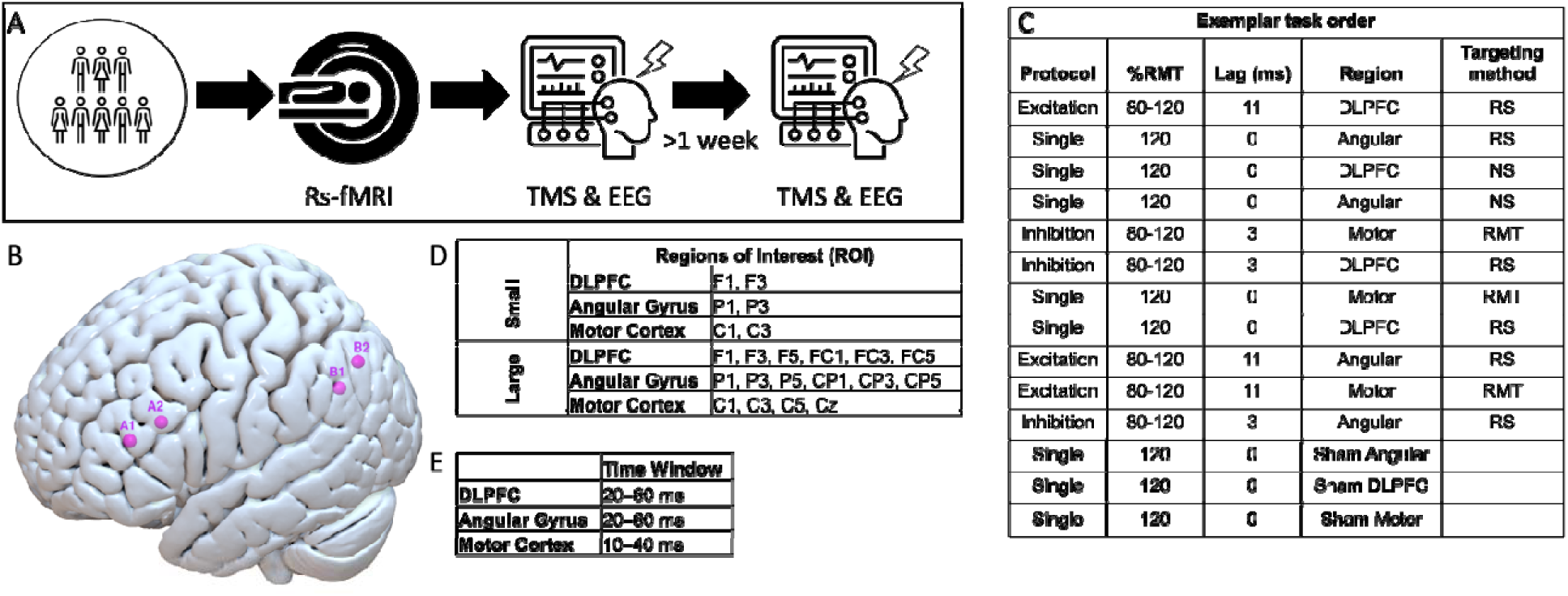
Study schematic. **A**) Study design: Following screening, participants underwent resting-state fMRI (rs-fMRI). Subsequently, they completed two TMS-EEG sessions, scheduled at least one week apart. **B**) Average individual connectivity- and Neurosynth-based targets for left DLPFC and left angular gyrus: A1: DLPFC_RS (−38, 39, 17), A2: DLPFC_NS (−38, 24, 24); B1: AngularGyrus_RS (−44, −58, 32), B2: AngularGyrus_NS (−46, −66, 44). **C)** Table depicts an exemplary sequence of experimental conditions for one subject. RMT: Resting Motor Threshold; RS: Resting State; NS: Neurosynth. **D)** Channels selected for small and large ROI definitions. Small ROI results are provided in the Supplemental Results. **E)** Time windows chosen for each region to capture of early neural responses in each target region, facilitating a comparison of TEP reliability across anatomically defined ROIs.

### TMS Setup

TMS was delivered using a Magstim-200 stimulator connected to a co-planar figure-of-eight Magstim-P/N9925 induction coil with a 70-mm wing radius (The Magstim Company Ltd., Whitland, UK). MRI-guided neuronavigation was performed using the Brainsight Neuronavigation system to ensure precise cortical stimulation and to minimize variability in the stimulation location across sessions.

### MRI Acquisition

Individual anatomical images required for 3D reconstruction and neuronavigation were obtained for each participant using a Philips Achieva 3T MRI scanner and a 32-channel head coil. A T1-weighted MPRAGE image was acquired for each participant, with a slice thickness of 1 mm, no slice gap, and a 256 x 256 matrix in sagittal orientation. Resting-state BOLD images were also collected to analyze intrinsic brain connectivity, with the following parameters: multi-slice (MS) gradient-echo planar imaging (EPI), Repetition Time (TR) of 800 ms, 910 volumes for a total scan time of 12:08, Echo Time (TE) of 30 ms, flip angle of 52°, slice thickness of 2.4 mm without slice gap, and in-plane resolution of 2.25mm^2^. Posterior to anterior and an anterior to posterior field maps were also acquired at the same resolution as the BOLD images to correct for distortion artifacts.

### MRI preprocessing

MRI data were organized using BIDS format^74^ and preprocessed using fMRIPrep 21.0.4^75,76^, which is based on Nipype 1.6.1^74,77^.

### Anatomical data preprocessing

The anatomical image for each participant was corrected for intensity non-uniformity (INU) using N4BiasFieldCorrection^78^, as distributed with ANTs 2.3.3^79^, and used as the anatomical reference throughout the workflow. The anatomical reference was then skull-stripped using a Nipype implementation of the antsBrainExtraction.sh workflow from ANTs, with OASIS30ANTs as the target template.

Brain tissue segmentation of cerebrospinal fluid (CSF), white matter (WM), and gray matter (GM) was performed on the skull-stripped anatomical image using fast [FSL 6.0.5.1]. Brain surfaces were reconstructed using recon-all [FreeSurfer 6.0.1^80^]. The brain mask estimated previously was refined with a custom method to reconcile ANTs-derived and FreeSurfer-derived segmentations of the cortical gray matter, as implemented in Mindboggle^81^.

Volume-based spatial normalization to a standard space (using the MNI152NLin2009cAsym template) was performed through nonlinear registration with antsRegistration (ANTs 2.3.3), using brain-extracted versions of both the T1w reference and the T1w template.

### Functional data preprocessing

For the BOLD run for each participant, a reference volume and its skull-stripped version were generated using a custom methodology of fMRIPrep. Head-motion parameters with respect to the BOLD reference (transformation matrices and six corresponding rotation and translation parameters) were estimated before any spatiotemporal filtering using mcflirt [FSL 6.0.5.1:57b01774^82^]. The BOLD time-series (including slice-timing correction) was then resampled to their original, native space by applying transforms to correct for head motion. These resampled BOLD time-series will be referred to as preprocessed BOLD in original space, or simply preprocessed BOLD.

The BOLD reference was then co-registered with six degrees of freedom to the anatomical reference using bbregister (FreeSurfer), which implements boundary-based registration^83^. Resampling was performed in a single interpolation step by composing all pertinent transformations (i.e., head-motion transform matrices, susceptibility distortion correction from fieldmaps, and co-registrations to anatomical and output spaces). Volumetric (gridded) resampling was performed using antsApplyTransforms (ANTs), configured with Lanczos interpolation to minimize smoothing effects from other kernels. Non-gridded (surface) resampling was performed using mri_vol2surf (FreeSurfer).

### rs-fMRI analysis

Individual resting-state scans were analyzed separately for each participant using Independent Component Analysis (ICA) to identify functional connectivity networks. Analysis was carried out using Probabilistic Independent Component Analysis^84^ as implemented in MELODIC (Multivariate Exploratory Linear Decomposition into Independent Components) Version 3.15, part of FSL (FMRIB’s Software Library, www.fmrib.ox.ac.uk/fsl).

The following pre-processing steps were applied to the input data: masking of non-brain voxels, voxel-wise demeaning of the data, and normalization of the voxel-wise variance. Pre-processed data were then whitened and projected into a 15-dimensional subspace using Principal Component Analysis.

The whitened observations were decomposed into sets of vectors that describe signal variation across the temporal domain (time-courses) and spatial domain (maps) by optimizing for non-Gaussian spatial source distributions using a fixed-point iteration technique^85^. Estimated component maps were divided by the standard deviation of the residual noise and thresholded by fitting a mixture model to the histogram of intensity values^84^.

### Target Selection

fMRI analysis for each participant was completed before they arrived for their first TMS session. Two fMRI-based independent components, one that comprised the left FPN and one that comprised the posterior DMN, were visually identified from the output of ICA. Peak coordinates for the FPN and DMN were selected for each participant, targeting the left DLPFC and left angular gyrus, respectively. The average connectivity-based DLPFC MNI coordinate across subjects was X=-38, Y=39, Z=17. The average connectivity-based angular gyrus MNI coordinate across subjects was X=-44, Y=-58, Z=32. Due to technical difficulties, the connectivity-based DLPFC session data for one participant and the angular gyrus excitation protocol session data for another participant were not saved, leaving 16 full datasets in those conditions.

For standardized target selection, peak voxels from a Neurosynth meta-analysis were used to locate the left DLPFC and left angular gyrus targets, based on the highest z-scores. MNI coordinates for Neurosynth-based DLPFC was X=-38, Y=24, Z=24. MNI coordinates for Neurosynth-based angular gyrus was X=-46, Y=-66, Z=44. Although literature-driven coordinates were considered, the consensus on angular gyrus coordinates in TMS studies was limited, so we opted for the meta-analysis approach instead.

The average Euclidean distance between the DLPFC definitions was 16.55 mm across subjects with the individualized DLPFC targets more anterior on average to the Neurosynth based target definition (**Figure 1B**). The average Euclidean distance between the angular gyrus definitions was 14.56 mm across subjects with the individual connectivity based angular gyrus targets being more anterior on average to the Neurosynth based target definition. The target for motor cortex stimulation was identified during identification of resting motor threshold at the beginning of each TMS session, following field-wide standards.

### Resting Motor Threshold (RMT)

The location of motor cortex stimulation was determined first using iterative movement of the TMS coil and stimulation to locate the coordinates that stimulate the abductor pollicis brevis (APB) muscle representation in the left motor cortex. Once found, the resting motor threshold (RMT) at this site was identified using electromyography with a descending relative frequency approach to identify the threshold at which an MEP of at least 50 μV is achieved in at least 50% of 10 to 20 consecutive trials^86,87^.

### TMS Stimulation Protocols

Motor cortex and each individual connectivity-defined target was stimulated using single-pulse and paired-pulse protocols. Neurosynth-defined targets were only stimulated using single-pulse protocol due to limitations on time and participant burden. The TMS coil was always held at an approximately 45 degrees angle relative to the midline. Stimulation intensity was set at 120% of the RMT for single-pulse TMS and at 80% (for the conditioning pulse) and 120% (for the test pulse) RMT for paired-pulse protocols. TMS was targeted for each region using MNI coordinates for each participant projected back into native space and then displayed on the participant’s anatomical image during neuronavigation. Sixty pulses were applied per protocol at each target site, with interstimulus intervals jittered around 5 seconds. Inter-session intervals ranged between 2 and 5 minutes.

The order of stimulation sites and intensities remained the same for each participant across sessions but was pseudorandomized between participants (see **Figure 1C** for an example sequence). This ensured that consecutive blocks were never over the same brain region, but any potential order effects remained constant within a participant. To help with data preprocessing, for the second half of the participants, at the end of the active TMS blocks, we administered one block of sham TMS over each brain region by positioning the coil near the participant’s head, maintaining contact without delivering actual stimulation by turning the coil 90-degree angle to the scalp. The intensity of the sham blocks was 120% of the RMT.

### TMS Setup

Neuronavigation ensured consistent coil positioning and orientation across both measurement sessions. The coil was manually held throughout the sessions and the distance error from the target per target region across subjects and sessions were as the following: motor cortex: 0.22 cm (SE=0.02) DLPFC_RS: 0.23 cm (SE=0.02) ; DLPFC_NS: 0.27 cm (SE=0.03) ; AngularGyrus_RS: 0.29 cm (SE=0.3); AngularGyrus_NS: 0.48 cm (SE=0.17).

For motor cortex stimulation, the coil was positioned to elicit maximal motor-evoked potentials (MEPs) during the first session, and this position was replicated in the second session. For the DLPFC, the coil was directed 45 degrees to the middle frontal gyrus, with the handle pointing laterally, as guided by the MRI-based neuronavigation system. For the angular gyrus, the coil was directed 45 degrees to the midline. The twist error from the target angle per target region across subjects and sessions were as the following: motor cortex: 3.28° (SE=1.07) DLPFC_RSl: 0.10° (SE=1.01) ; DLPFC_NS: 0.85° (SE=0.90) ; AngularGyrus_RS: 2.06° (SE=1.35) ; AngularGyrus_NS: 0.89° (SE=1.38).

### EEG Recording

EEG responses to TMS were recorded using a 64-channel TMS-compatible EEG system (ActiChamp, Brain Products, GmbH, Germany) with passive electrodes designed to prevent overheating due to eddy currents induced by TMS. The EEG system was connected to a TMS-compatible amplifier with a sampling rate of 25 kHz, a bandwidth of 0.1–350 Hz, and a 16-bit AD conversion resolution. The EEG array was configured according to the international 10-20 system for optimal scalp coverage. EasyCap Abralyt HiCl High chloride abrasive electrolyte gel was used. Impedances were monitored after each session and the channels had impedances lower than 5 kΩ, ensuring good signal quality. The reference channel used during recording was Fz; the ground channel used during recording was FCz.

### EEG Data Preprocessing

The EEG data were segmented around each TMS pulse, with epochs spanning from −1.5 to 1.5 seconds relative to the test pulse in paired-pulse protocols. Baseline correction was applied differently for each protocol. For single-pulse protocols, baseline correction was performed using the interval from −500 to −5 milliseconds before the pulse. For paired-pulse inhibition protocols, the baseline interval was adjusted to −500 to −8 milliseconds before the test pulse, and for excitation protocols, it was adjusted to −500 to −16 milliseconds before the test pulse.

Condition-specific artifact removal was performed to address TMS-related noise. For single-pulse protocols, data from −2 to 5 milliseconds were cut. For inhibition protocols, data from −5 to 5 milliseconds before the test pulse were cut, and for excitation protocols, from −13 to 5 milliseconds before the test pulse were cut. After this initial artifact removal, bad channels and trials were excluded to improve data quality. An average of 1.7 trials [0 11] (*SD*=3.90) and 0.77 channels [0 17] (*SD*=1.21) were rejected. The data were detrended and underwent baseline correction to remove any residual artifacts. Further processing included Independent

Component Analysis (ICA). Data were reduced to 30 components to extract and remove eye blink, muscle, and line noise artifacts. For the second half of the sample, sham blocks were acquired per region and were concatenated to the end of real stimulation data only during the ICA step separately for each stimulation site to help identify artifacts. The Source-estimate-utilizing noise-discarding algorithm (SOUND^88^) was applied to suppress extracranial noise. The Signal-space projection–source-informed reconstruction (SSP-SIR^89^) method was used to reduce muscle artifacts induced by TMS. After artifact suppression, the missing time intervals were interpolated, and the data were bandpass-filtered with cutoff frequencies of 1–100 Hz and bandstop-filtered with cutoff frequencies of 58–62 Hz to remove slow drifts, high-frequency noise and line noise.

### TEP Data Analysis

To determine the time course of TMS-evoked potentials (TEPs) for each target region, we defined both large and small regions of interest (ROIs) based on sets of channels covering the DLPFC, motor cortex, and angular gyrus. Per each region, small and large ROI definitions were chosen (see **Figure 1D**). Specific time windows (**Figure 1E**) were selected to optimize the capture of early neural responses in each target region, minimizing the influence of sensory confounds typically affecting later components. This approach facilitates the comparison of TEP reliability across anatomically defined ROIs.

We calculated the peak-to-peak amplitude of TEPs by extracting data from predefined channels above within the specified time windows. We computed the peak-to-peak amplitude by identifying the maximum and minimum values within the time window and calculating their difference. The resulting amplitude was then converted from linear microvolts (μV) to decibels (dB) using a logarithmic transformation, providing a consistent measure of TEP response strength across conditions.

To determine the test-retest reliability of early TEPs at different time points, we used the Concordance Correlation Coefficient (CCC), a measure optimized for assessing test-retest reliability^69,90^. The CCC compares the variance between subjects to the total variance to evaluate the agreement between two sets of observations. We applied the CCC to quantify the reliability of TEPs across different stimulation conditions by comparing the magnitude of TEPs from one session to another session. The CCC values were calculated for each target and time point, with mean values and 95% confidence intervals reported. To compare CCC values across conditions, we employed the percentile bootstrap method^91^ to assess statistical differences in reliability^68^.

## Results

In this study, we assessed the test-retest reliability of early TEPs for functionally defined targets derived from the RMT procedure and fMRI. Specifically, we examined three left-lateralized brain regions—motor cortex (M1), dorsolateral prefrontal cortex (DLPFC), and angular gyrus—using three TMS protocols: single-pulse TMS, short-interval intracortical inhibition (SICI), and long-interval intracortical inhibition (LICI). For motor cortex, targets were defined using a standard RMT hotspotting procedure. For DLPFC and angular gyrus, targets were defined using two methods: individual resting-state (RS) fMRI-based target localization and functional definition based on Neurosynth coordinates on normalized anatomical images. TEP reliability was assessed using the concordance correlation coefficient (CCC) for within-session and between-session reliability of the logarithmic peak-to-peak amplitude quantification within large ROIs, with bootstrapped confidence intervals (CIs) calculated for each CCC estimate (see Supplemental Results for smaller ROI definitions and linear values). See Figures 1, 2, and 3 for the time course (**figures 2 and 4**) and scalp distribution (**figure 3**) of TEPs across all participants for each experimental condition.

**Figure 2.**
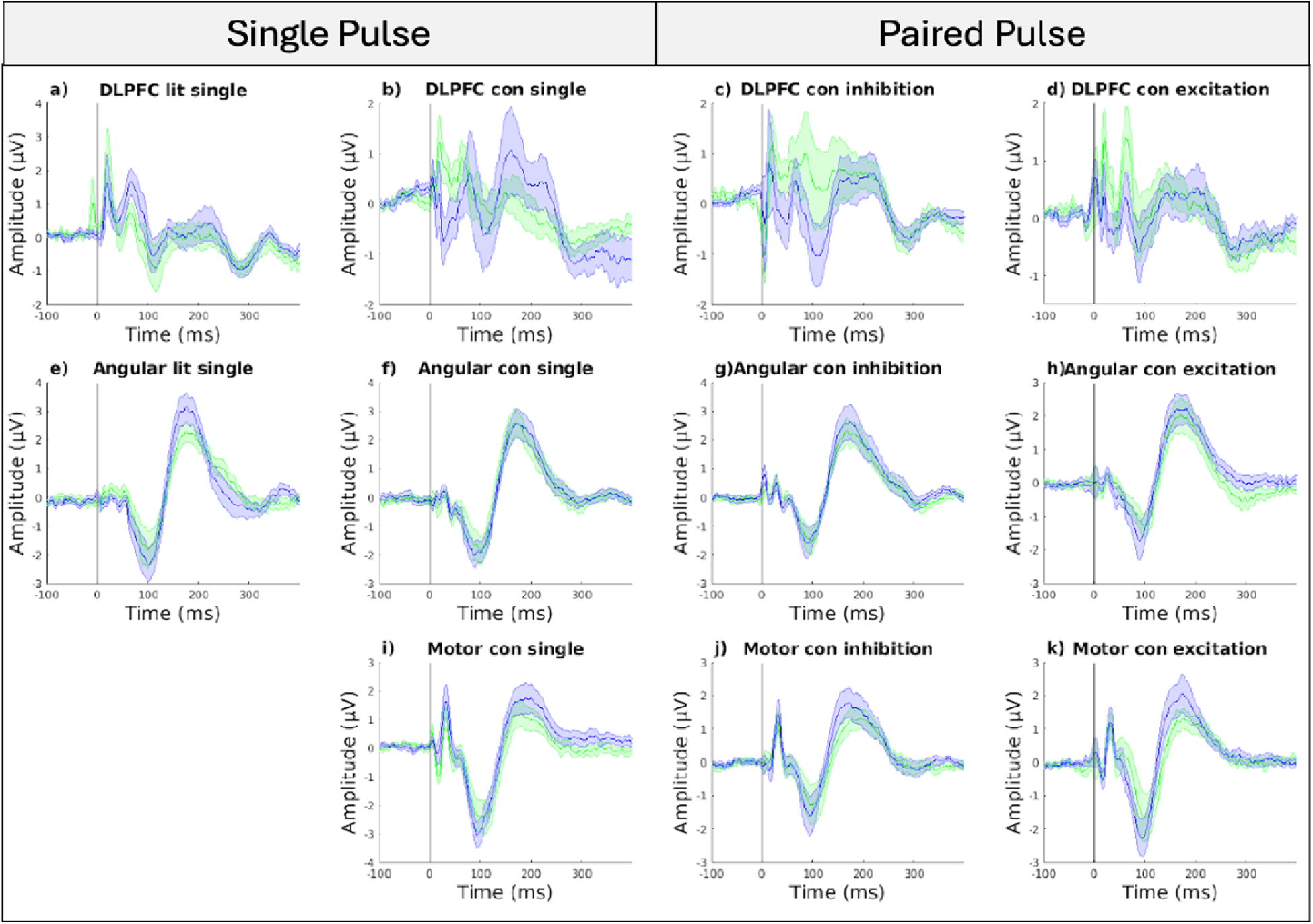
Time course of TEPs across all participants for each experimental condition. Green lines stand for session 1 values; blue lines stand for session 2 values. Time is shown relative to the test pulse in excitation and inhibition conditions. RS: Resting State; NS: Neurosynth.

**Figure 3.**
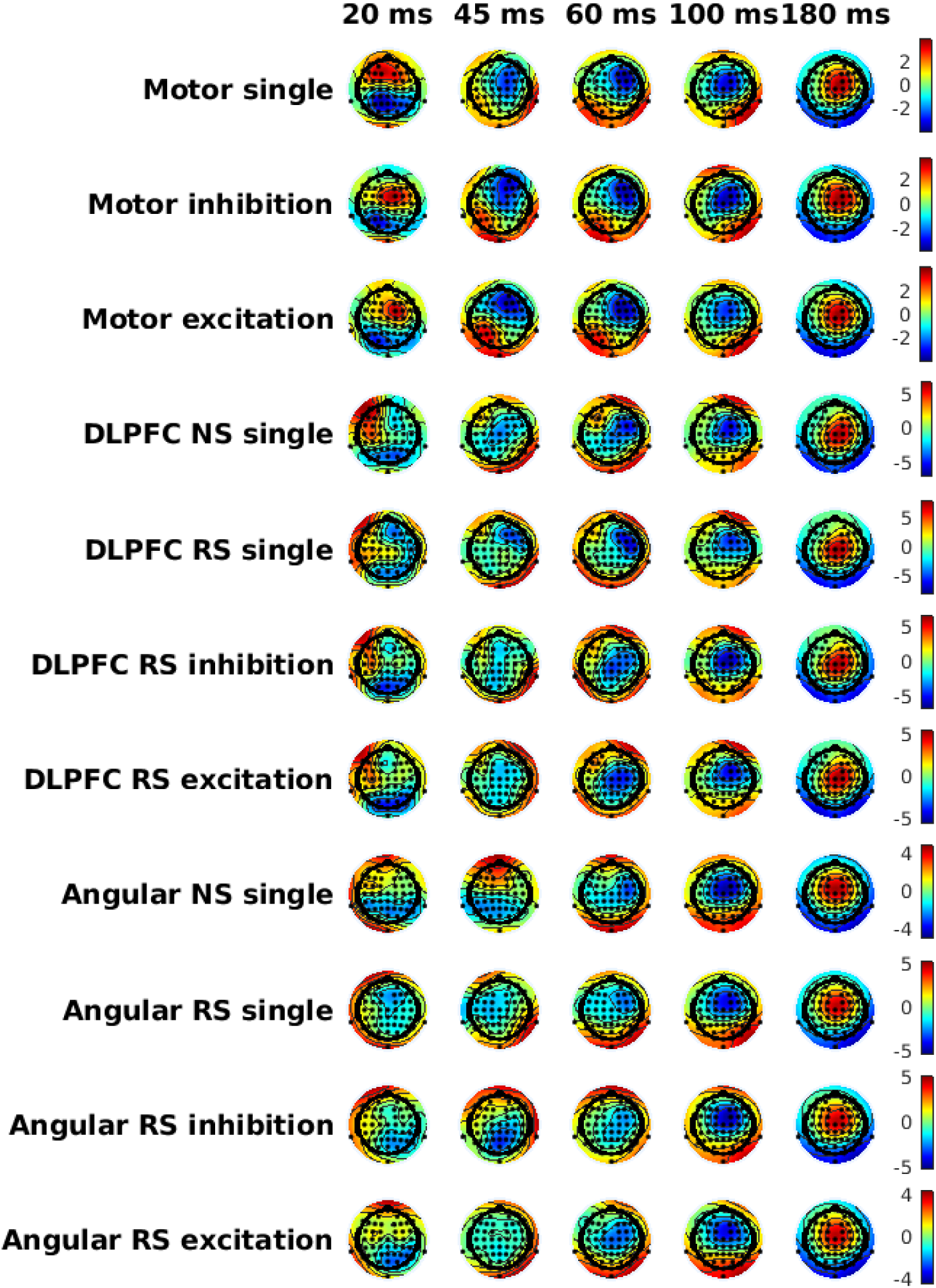
Topographical maps across all participants and sessions for each experimental condition, averaged across sessions. Time is shown relative to the test pulse in excitation and inhibition conditions. RS: Resting State; NS: Neurosynth.

**Figure 4.**
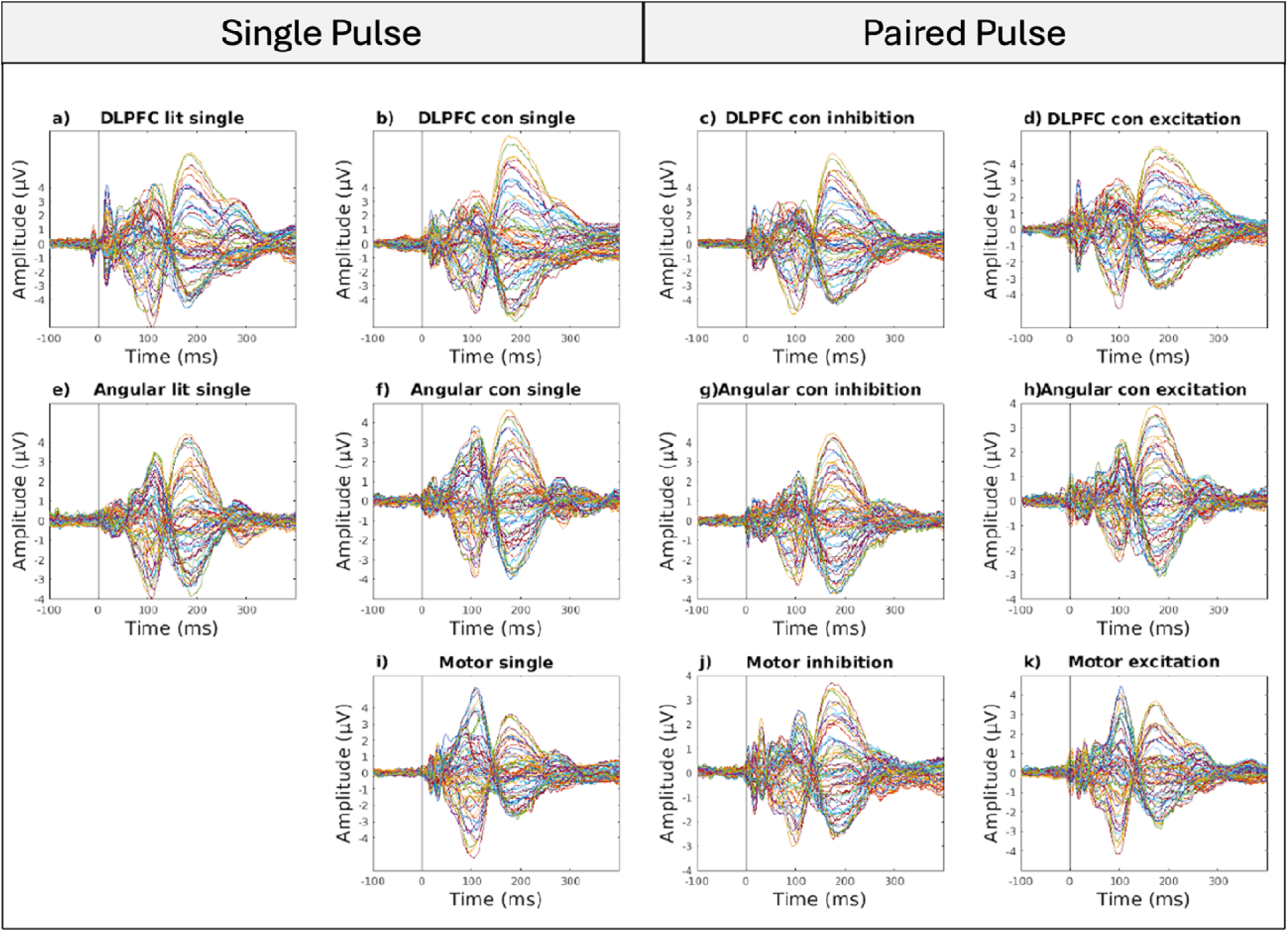
Butterfly plots across all participants and sessions for each experimental condition averaged across sessions. Colored lines stand for channels. Time is shown relative to the test pulse in excitation and inhibition conditions. RS: Resting State; NS: Neurosynth.

### Reliability Across Brain Regions

TEP reliability varied across the three regions of interest (ROIs), with the highest CCC values observed in the motor cortex (CCC_mean_ = 0.59, CCC_std_ = 0.08) and lowest reliability scores for DLPFC (CCC_mean_ = 0.40, CCC_std_ = 0.15) and angular gyrus between them (CCC_mean_ = 0.45, CCC_std_ = 0.20) (**Figure 5**). These results indicated that motor cortex stimulation overall produced the most consistent TEPs across regions.

**Figure 5.**
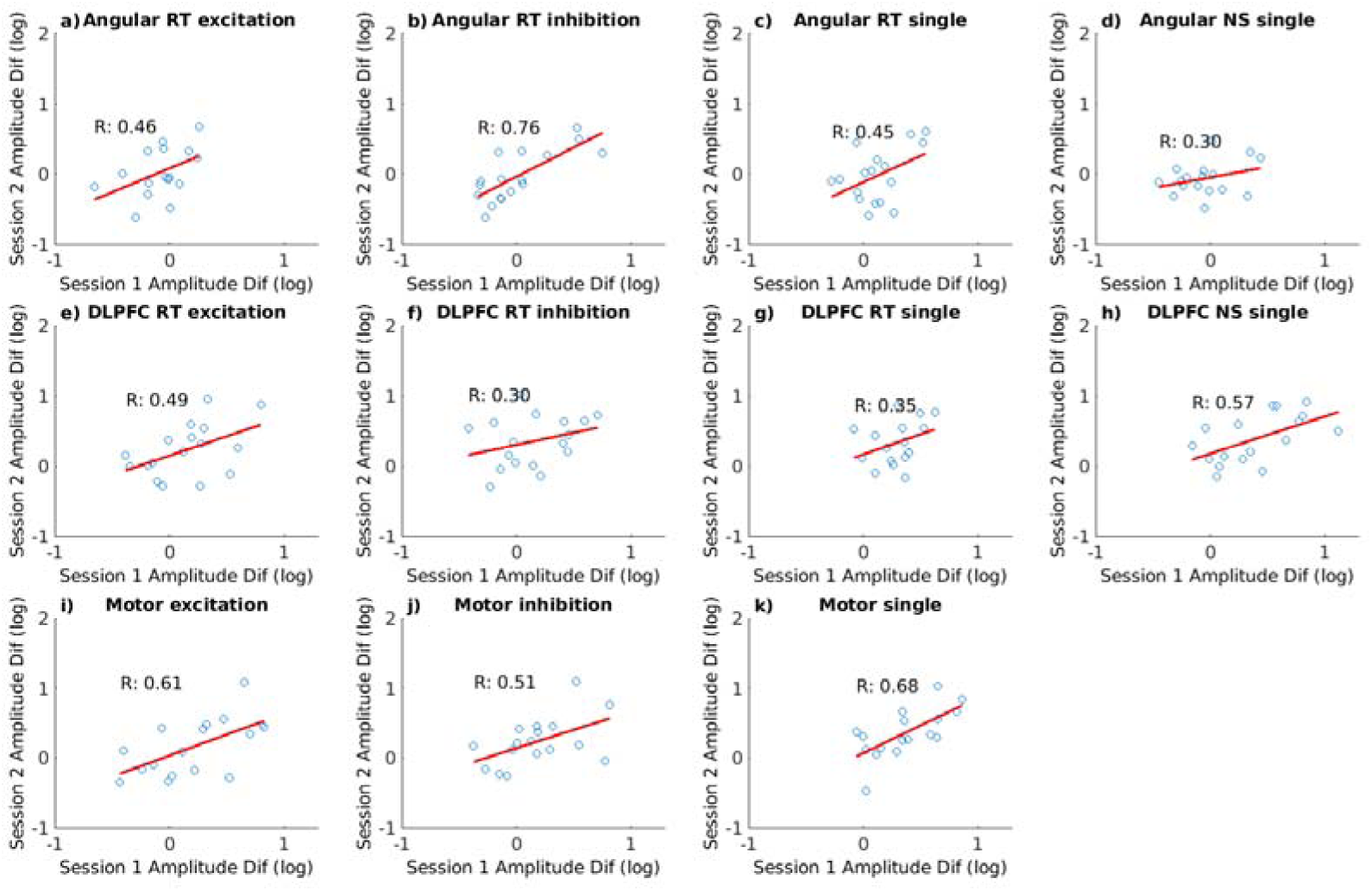
Scatter plot of log-transformed amplitude differences, calculated by taking the difference between maximum and minimum peak-to-peak amplitudes within a time window, between sessions across experimental conditions. R values stand for Pearson product moment correlation coefficients. RS: Resting State; NS: Neurosynth.

### Reliability Across Target Definitions

For DLPFC and angular gyrus, we compared reliability for targets defined using RS and Neurosynth-derived functional overlays in the single pulse condition. Reliability for each experimental condition is presented in **Table 1**. Neurosynth-defined targets generally yielded higher CCC values in DLPFC with a mean CCC range of 0.57, CI [0.14 0.82], compared to RS derived targets, which showed a CCC range of 0.30, CI [-0.14 0.64] across sessions. However, the CCC difference between different DLPFC definitions was not significant, as revealed by bootstrap statistics (CI_difference_ [-0.87 0.01], *P* = 0.06).

**Table 1.** Reliability of TEPs between session 1 and session 2, across experimental condition. CCC: concordance correlation coefficient; CI: 95% confidence interval; Mean Amp: mean peak-to-peak amplitude of the signal within selected time windows; RS: Resting State; NS: Neurosynth.

|  | CCC | low CI | high CI | Mean Amp |
| --- | --- | --- | --- | --- |
| Angular RS excitation | 0.40 | -0.03 | 0.71 | -0.03 |
| Angular RS inhibition | 0.75 | 0.44 | 0.90 | -0.01 |
| Angular RS single | 0.35 | -0.03 | 0.64 | 0.05 |
| Angular NS single | 0.30 | -0.19 | 0.67 | -0.05 |
| DLPFC RS excitation | 0.47 | 0.02 | 0.76 | 0.18 |
| DLPFC RS inhibition | 0.25 | -0.17 | 0.59 | 0.25 |
| DLPFC RS single | 0.30 | -0.14 | 0.64 | 0.31 |
| DLPFC NS single | 0.57 | 0.14 | 0.82 | 0.39 |
| **Motor excitation** | 0.60 | 0.19 | 0.83 | 0.19 |
| **Motor inhibition** | 0.51 | 0.06 | 0.78 | 0.21 |
| **Motor single** | 0.67 | 0.31 | 0.86 | 0.36 |

In angular gyrus, both target definitions showed moderate reliability with overlapping confidence intervals (Neurosynth-based: CCC = 0.30, CI [-0.19 0.67]; RS: CCC = 0.35, CI [-0.03 0.64]. The CCC difference between different angular gyrus definitions was not significant, as revealed by bootstrap statistics (CI_difference_ [-0.55 0.70], *P* = 0.83).

### Effect of TMS Protocol on TEP Reliability

There was no statistical difference between single-pulse and SICI (CI_difference_ [-0.26 0.43], *P* = 0.69) or LICI (CI_difference_ [-0.24 0.63], *P* = 0.40) conditions for M1. Similarly, there was no statistical difference between single-pulse and SICI (CI_difference_ [-0.65 0.36], *P* = 0.53) or LICI (CI_difference_ [-0.55 0.52], *P* = 0.91) conditions for DLPFC. There was also no statistical difference between single-pulse and SICI (CI_difference_ [-0.52 0.43], *P* = 0.93) or LICI (CI_difference_ [-0.86 0.04], *P* = 0.08) conditions for angular gyrus. These results indicate that all three TMS protocols have similar reliability across regions.

## Discussion

This study assessed the test-retest reliability of early TEPs in the DLPFC and angular gyrus, using the motor cortex (M1) as a benchmark. Three TMS protocols (single-pulse, SICI, and LICI) and two targeting methods (RS- and NS-based) for DLPFC and angular gyrus were evaluated. While Neurosynth-derived targets showed slightly higher reliability for DLPFC compared to resting-state targets, angular gyrus results were similar across targeting methods. Across all regions, no significant reliability differences were observed between the three protocols. Notably, individualized functional neuroimaging targets for DLPFC and angular gyrus did not improve TEP reliability over the RMT-based method for M1.

Central targets like M1 generally exhibit greater TEP reliability in TMS-EEG studies than lateral regions such as DLPFC and angular gyrus due to reduced susceptibility to muscle artifacts^63^. Muscle activation in lateral targets, such as the frontalis muscle for DLPFC and temporalis muscle for angular gyrus, causes artifacts sensitive to coil positioning and intensity^62,64^. While adjustments like coil orientation, reduced stimulation intensity, and offline artifact removal (e.g., ICA) can mitigate these effects, lateral targets remain more artifact-prone. In contrast, M1 benefits from cleaner EEG signals and real-time optimization using EMG feedback. Extending such real-time feedback to non-motor regions may enhance TEP reliability by allowing immediate adjustments to site, orientation, and intensity^92,93,94^.

Our findings align with Gogulski et al.^68^, who reported that anterior DLPFC targets yield smaller TEP amplitudes and lower reliability compared to posterior targets. Smaller amplitudes may reflect reduced neural engagement or increased artifact susceptibility at anterior sites. Similarly, our study observed numerically lower TEP reliability for anterior connectivity-based DLPFC targets compared to Neurosynth-derived targets. This highlights the need for careful validation of functionally guided targets to optimize amplitude and reliability.

Kerwin et al.^38^ emphasized the influence of trial number, target location, and interval duration on TMS-EEG reliability. In our study, 60 trials per condition over a one-week interval yielded a CCC of 0.57 for DLPFC targets, lower than Kerwin et al.’s >0.8 concordance for peaks such as N100 and P200 using shorter intervals. Longer intervals and non-central targets likely contributed to reduced reliability, consistent with prior findings^36^. Gogulski et al.^68^ examined the number of TMS pulses required for reliable prefrontal EL-TEPs, finding that 50 trials were sufficient for most targets, except for the most anterior dlPFC target, which showed low reliability regardless of trial count. Notably, the medial target achieved high reliability with as few as 25 trials across within-block, between-block, and test-retest assessments, indicating that for targets low in reliability, increasing the number of trials may not improve consistency. Increasing trial numbers and refining targeting approaches may improve reliability in future studies.

Stimulation intensity also affects TEP reliability. While we used standardized intensities (120% and 80% RMT), Mutanen et al.^63^ showed that reducing intensity to 60–80% of RMT substantially decreases muscle artifacts, a major noise source in TEP recordings. On the other hand, in M1, the supra-threshold intensity likely elicited ascending spinal-to-cortical input^95^, potentially confounding the TEPs and contributing to the higher CCCs observed. This suggests that optimal intensity settings for TEP studies may differ from clinical practices.

Our results underscore differences between TMS-fMRI and TMS-EEG targeting. While fMRI-based approaches prioritize functional connectivity to engage networks (e.g., DLPFC-subgenual cingulate in depression), TMS-EEG must address artifacts that obscure neural signals^94^.

Identifying targets with the highest signal-to-noise ratio (SNR) may be more effective for TMS-EEG studies than strictly following fMRI connectivity targets.

Integrative approaches, such as combined TMS-fMRI, tractography, and high-density EEG, offer a path forward^96^. These methods could help to optimize both target engagement and TEP reliability by mapping structural, functional, and connectional profiles of stimulation sites. By aligning TMS-fMRI’s network engagement with TMS-EEG’s focus on SNR, future studies could advance the reliability of TMS-EEG biomarkers while maintaining individualized targeting precision.

## Acknowledgements

This work was supported by the Johns Hopkins Center for Psychedelic and Consciousness Research, which was funded by a generous gift from the Steven and Alexandra Cohen Foundation as well as Tim Ferriss, Matt Mullenweg, Blake Mycoskie, and Craig Nerenberg.

We thank Daniel Jaro, Eli Weisman, and Gabi Lofland for their contributions to the preprocessing of TMS/EEG data. Chris Cline for providing scripts for data visualization. We also thank Daniel Galvez Cabrera and Sean Garcy for their assistance with data collection.

## Availability of data and materials

Data is available to share upon request.

## Code availability

The authors will upload all relevant analysis code on a public repository.

## Authors’ contributions

CS and FB conceptualized the work. FB provided the funding for the study. CS collected and analyzed the data. JG, IG and PL provided support for data analysis and interpretation of data. All authors wrote, revised, and approved of the final manuscript.

## Consent to participate

All human subjects provided informed consent to participate in this study. <u>Consent for publication:</u> All human subjects provided informed consent to publish this research.

## Supplementary Results

**Table S1.**
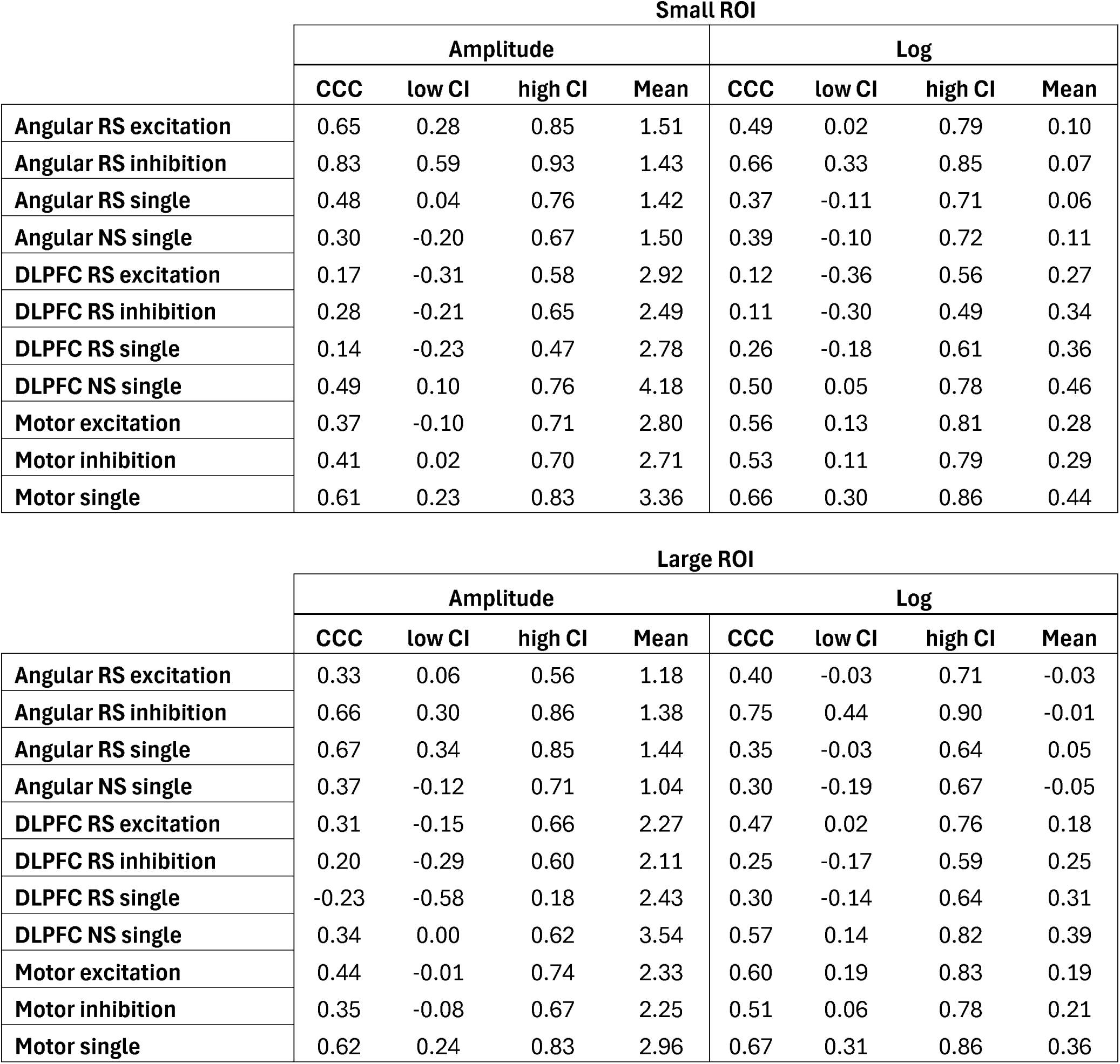
The table shows the CCCs across experimental conditions with small and large ROIs and linear and logarithmic peak-to-peak amplitudes with 95% CI.

|  | Small ROI |  |  |  |  |  |  |  |
| --- | --- | --- | --- | --- | --- | --- | --- | --- |
|  | Amplitude |  |  |  | Log |  |  |  |
|  | CCC | low CI | high CI | Mean | CCC | low CI | high CI | Mean |
| Angular RS excitation | 0.65 | 0.28 | 0.85 | 1.51 | 0.49 | 0.02 | 0.79 | 0.10 |
| Angular RS inhibition | 0.83 | 0.59 | 0.93 | 1.43 | 0.66 | 0.33 | 0.85 | 0.07 |
| Angular RS single | 0.48 | 0.04 | 0.76 | 1.42 | 0.37 | -0.11 | 0.71 | 0.06 |
| Angular NS single | 0.30 | -0.20 | 0.67 | 1.50 | 0.39 | -0.10 | 0.72 | 0.11 |
| DLPFC RS excitation | 0.17 | -0.31 | 0.58 | 2.92 | 0.12 | -0.36 | 0.56 | 0.27 |
| DLPFC RS inhibition | 0.28 | -0.21 | 0.65 | 2.49 | 0.11 | -0.30 | 0.49 | 0.34 |
| DLPFC RS single | 0.14 | -0.23 | 0.47 | 2.78 | 0.26 | -0.18 | 0.61 | 0.36 |
| DLPFC NS single | 0.49 | 0.10 | 0.76 | 4.18 | 0.50 | 0.05 | 0.78 | 0.46 |
| Motor excitation | 0.37 | -0.10 | 0.71 | 2.80 | 0.56 | 0.13 | 0.81 | 0.28 |
| Motor inhibition | 0.41 | 0.02 | 0.70 | 2.71 | 0.53 | 0.11 | 0.79 | 0.29 |
| Motor single | 0.61 | 0.23 | 0.83 | 3.36 | 0.66 | 0.30 | 0.86 | 0.44 |

|  | Large ROI |  |  |  |  |  |  |  |
| --- | --- | --- | --- | --- | --- | --- | --- | --- |
|  | Amplitude |  |  |  | Log |  |  |  |
|  | CCC | low CI | high CI | Mean | CCC | low CI | high CI | Mean |
| Angular RS excitation | 0.33 | 0.06 | 0.56 | 1.18 | 0.40 | -0.03 | 0.71 | -0.03 |
| Angular RS inhibition | 0.66 | 0.30 | 0.86 | 1.38 | 0.75 | 0.44 | 0.90 | -0.01 |
| Angular RS single | 0.67 | 0.34 | 0.85 | 1.44 | 0.35 | -0.03 | 0.64 | 0.05 |
| Angular NS single | 0.37 | -0.12 | 0.71 | 1.04 | 0.30 | -0.19 | 0.67 | -0.05 |
| DLPFC RS excitation | 0.31 | -0.15 | 0.66 | 2.27 | 0.47 | 0.02 | 0.76 | 0.18 |
| DLPFC RS inhibition | 0.20 | -0.29 | 0.60 | 2.11 | 0.25 | -0.17 | 0.59 | 0.25 |
| DLPFC RS single | -0.23 | -0.58 | 0.18 | 2.43 | 0.30 | -0.14 | 0.64 | 0.31 |
| DLPFC NS single | 0.34 | 0.00 | 0.62 | 3.54 | 0.57 | 0.14 | 0.82 | 0.39 |
| Motor excitation | 0.44 | -0.01 | 0.74 | 2.33 | 0.60 | 0.19 | 0.83 | 0.19 |
| Motor inhibition | 0.35 | -0.08 | 0.67 | 2.25 | 0.51 | 0.06 | 0.78 | 0.21 |
| Motor single | 0.62 | 0.24 | 0.83 | 2.96 | 0.67 | 0.31 | 0.86 | 0.36 |

